# Retro Nasal blockade reduces the Neural Processing of Sucrose in the Human Brain

**DOI:** 10.1101/2025.02.11.637706

**Authors:** Hee-kyoung Ko, Jingang Shi, Thomas Eidenberger, Weiyao Shi, Ciara McCabe

## Abstract

It is assumed “Non-volatile” tastes like sucrose do not activate retro nasal pathways. Recent studies find that sucrose when aerosolized, can reach the retro nasal olfactory region and be perceived. The neural mechanisms by which the human brain interprets sucrose via retro nasal pathways is unknown.

We examined neural activity to sucrose with a nose clip on (blocking retro nasal) and nose clip off, in healthy adults (N=34, mean 25 yrs.). We examined the whole brain and ROIs involved in taste, smell, attention, reward and multi-modal integration; insula, postcentral gyrus, amygdala, olfactory cortex, subgenual and pregenual anterior cingulate, nucleus accumbens and OFC. We also examined correlations with subjective ratings of pleasantness and mouth fullness. We also examined the effect of the nose clip on the time to peak activity for sucrose using the bold signal time course.

The nose clip on vs off reduced the subjective experience of mouth fullness. Neural activity to sucrose was reduced with the nose clip on in the primary taste, olfactory, attention and reward ROIs and in the rolandic operculum, lingual gyrus and precuneus in the whole brain analyses. The olfactory and prefrontal cortex ROIs tracked subjective mouth fullness, but this was not apparent with the nose clip on.

Blocking retro nasal sensation reduces subjective and neural responses to sucrose taste. Retro nasal sensations could play a role in “pure” taste perception. Developing more satisfying low-sugar foods could be achieved by enhancing the perception of sweetness through aroma modulation.

## Introduction

Volatile odour molecules released from food or drink in the mouth travel up the back of the throat into the nasal cavity and activate the olfactory receptors via retro nasal pathways (Buettner et al., 2001; Stevens & Cain, 1986; Voirol & Daget, 1986). Non-volatile taste compounds such as sucrose however are thought recognised by the gustatory system (Roper & Chaudhari, 2017) via purely oral pathways. As such, tastes like sucrose that are considered non-volatile should not activate retro nasal pathways.

Yet early work has shown retro nasal sensation does play a role in how humans experience non-volatile tastes such as sucrose. Disabling retro nasal sensation through reversed nasal airflow significantly impaired participants’ ability to identify sucrose, although most were still able to perceive its sweetness (Mozell et al., 1969). Similarly studies have reported that blocking retro nasal sensation with a nose clip notably increases detection and recognition thresholds (Murphy & Cain, 1980), reduces identification accuracy (Masaoka et al., 2010), and diminishes the perceived sweetness intensity of sucrose solutions (Mojet et al., 2003; Mu et al., 2024; Yang et al., 2024). As expected, Yang et al. found that the nose clip only effected the taste and olfactory senses and not other senses such as vision or touch, which suggests an important component of olfactory processing even with “non-volatile” tastes (Yang et al., 2024).

Studies also report that the observed differences in sucrose sweetness perception in young people vs old diminishes when youth are wearing a nose clip, suggesting retro nasal sensations account for the differences (Mojet et al., 2005). Furthermore, studies find that nasal blockage via sinusitis can also significantly reduce sucrose detection (Tsuji et al., 2018). Thus it is possible that everyday colds could affect taste perception via retro nasal blockade (James et al., 2022) and viruses such as COVID19 could also impact taste perception via retro nasal dysfunction. As this can have serious psychological implications in the longer term (Javed et al., 2022) it is imperative to understand the contribution of retro nasal pathways to taste processing.

Given sucrose is considered non-volatile such studies have led to the idea that it is impurities in taste solutions rather than the tastants themselves that is being recognised by retro nasal processing (Mojet et al., 2005). Others refute this hypothesis as they observed that the purity grade of the tastants (e.g., sucrose in reagent grade, non-reagent grade, and food grade) did not influence olfactory discrimination ability, both in mice (Zukerman et al., 2009) and in humans (Chen, 2013).

One explanation for reduced perception with nasal blockage could be that non-volatile compounds such as sucrose taste do indeed activate the retro nasal pathways. Our recent study using high speed cameras found that an orally-ingested sucrose solution could be transferred to the nasal cavity in the form of aerosol particles (He et al., 2023). This plausibly explains why and how retro nasal sensation is involved in the oral consumption of non-volatile sucrose, affecting its identification, intensity perception and threshold detection. As sucrose sweetness intensity was reduced when the volunteers’ noses were clipped, this also indicates the involvement of retro nasal sensation during its drinking (He et al., 2023). These findings were extended with our findings that retro nasal sensation can contribute to the discrimination between tastes such as sucrose and sucralose and to the perception of sweeteners (He et al., 2024).

While these findings clearly highlight the involvement of retro nasal sensation in the perception of sucrose, the underlying neural mechanisms underpinning the involvement of the retro nasal pathway vs the ortho nasal pathway in sucrose taste perception is unknown.

Taste processing is first achieved at the level of taste receptor cells which are clustered in taste buds on the tongue. When receptors are activated by specific tastants, they transmit information via sensory afferent fibres to specific areas in the brain that are involved in taste perception (Lee & Owyang, 2019). Functional magnetic resonance imaging (fMRI) reveals that taste activates the anterior insula/frontal operculum, the primary taste cortex, the orbitofrontal cortex (OFC) (possibly secondary taste cortex) and the anterior cingulate (ACC) (Rolls, 2019). It’s also been shown that the intensity of tastes is correlated with activity in the insula whereas pleasantness is in prefrontal regions like the orbitofrontal cortex and anterior cingulate (Rolls, 2019).

Retro nasal processing allows molecules to reach the olfactory epithelium in the nasal cavity, where they bind to specific olfactory receptors. The receptors send electrical signals via the olfactory nerve to the olfactory bulb, located at the base of the brain. From here the signal is relayed to higher brain areas such as the piriform cortex the primary olfactory area for odour perception. In humans, the piriform cortex is correlated with the intensity of odours but not their pleasantness (Rolls, 2019). Signals from the olfactory bulb also project to the orbitofrontal cortex (OFC) where odour and other visual and sensory information are combined to contribute to stimuli identification and evaluation (Rolls, 2015, 2019). However, if non-volatile taste compounds such as sucrose are also perceived through retro nasal pathways, it raises the question of whether the blockade of retro nasal sensation would reduce or slow down the integration of neural responses to sucrose.

Sucrose is known not only for its sweetness but also for contributing to a sensation of fullness or mouthfeel (Lavin et al., 2002). Studies have indicated that nasal occlusion can diminish the perception of fullness (Baraniuk, 2011; Yeomans & Boakes, 2016) which given our recent research on sucrose and retro nasal processing could indicate a roll for retro nasal sensation in not only sweetness but fullness too. Further, the neural activity underpinning sucrose pleasantness and fullness in relation to retro nasal sensation remains unexplored.

Therefore, the aim of this study was to map the brain activity to sucrose taste activated by oral cavity (nose-clip-on, blocking retro nasal contribution) and the contribution of retro nasal cavity activation (nose-clip-off). We could do this by delivering sucrose taste to the mouth of the participant, whilst they are in the MRI machine. We examined the effects of sucrose vs a control taste on whole brain activity and examined regions of interest such as the primary taste and olfactory areas and multi modal integrations areas like the OFC. We also examined regions such as the amygdala which responds to taste and oral texture (Kadohisa et al., 2005) and which projects to olfactory regions and responds to both pleasant and unpleasant tastes (O’Doherty et al., 2001a; Pu et al., 2024), the perigenual cingulate (pgACC) highlighted in a study examining differences in neural responses to ortho nasal vs retro nasal odour delivery (Small et al., 2005). We also examined the subgenual ACC (sgACC) an area thought to involuntarily attend to odours (Veldhuizen & Small, 2011). Finally, as we are interested in the retro nasal contribution to the rewarding effects of sucrose we also examined the nucleus accumbens (Berridge, 2009). We extracted activity in the ROIs and examined activity between nose clip off vs on conditions to assess the contribution of retro nasal sensation to neural activity. We also examined the correlation between the ROI activity and subjective ratings of pleasantness and mouth fullness taken during the scanning. As we are also interested in the effects of retro nasal processing on the timing of activity, we also examined the time to peak activity within 10 s, in the nose clip on vs off conditions. This was examined by calculating the time to maximum peak activity within the first 10 s from the onset of stim delivery from the time course data, using the max function in Matlab.

## Materials and Methods

### Participants

Thirty-four healthy and right-handed adults (10 male and 24 female) were recruited for the fMRI study. All participants were between 18 and 45 years old (mean 25.71 ± 8.25) and had a current body mass index (BMI, weight in kg/height in m2) or waist-to-height ratio (WTH) in the healthy range. Participants were excluded if they had any current/previous psychiatric history using the Structured Clinical Interview for DSM-IV Axis I Disorder Schedule (SCID), or if they took psychoactive medication or an eating disorder (measured with Eating Attitude Test (EAT) > 20), food allergies, diabetes, smoking, or any contraindications to fMRI scanning. We also recorded the frequency, liking and craving for sugary and sweetened foods (Rolls & McCabe, 2007). The questions in this scale consisted of “How frequently do you eat sugary foods?” with answers of either; a few times per month; 1-2 times per week; 3-4 times per week; or more than 5 times per week and “How frequently do you eat/drink foods with sweeteners?”, with answers of either; Never; Rarely; Sometimes; Often; Usually or Always. The Craving and Liking for sugary foods were scored as 1 for low craving and 10 for high craving on a Likert scale. All procedures contributing to this work comply with the ethical standards of the Helsinki Declaration of 1975, as revised in 2013 and ethical approval was obtained from the University of Reading Ethics committee, ethics ref: 2023-130-CM all participants provided written informed consent.

### Pre-test 1 (Triangle test or Taste perception test)

The 34 participants were entered into the study if they could distinguish 2% sucrose from a control. This standard taste perception test was as follows: The participants were randomly allocated to the following sequences of two samples A (distilled water) and B (20 g sucrose/litre [2 % Sucrose]): ABB, AAB, ABA, BBA, BAA and BAB. For the individual performance, each participant received all six sequences in random order. In a sequence, the participants took the whole 10 mL of each sample into their mouth, swirled and coat the solution around their mouth for 3 seconds and then spit it into a spittoon. On each trial after tasting all three, they indicated which was different from the other two. Participants who yielded correct identification of at least 5 out of the 6 trials on a second attempt, were recruited to the study.

### Pre-test 2 (Candy smell test retro nasal)

We used the candy smell test to check participants retro nasal olfactory performance (Renner et al., 2009). This test examines participants ability to identify the flavour of a candy (500 mg) placed on the middle of the tongue from 6 possible choices (6-alternative, forced-choice procedure) strawberry, banana, orange, coffee, cherry, or pineapple. Participants can suck the candy or chew it if necessary. The participants wrote down one of the choices. If they cannot identify the candy they can skip to the next trial. Each participant performed 5 trials with nose clip on and 5 trials with nose clip off. Between trials participants rinse their mouths with water. There was no feedback to the participants whether their responses were correct or incorrect. We expected less than 40 - 50% correct (1 or 2 corrects / 5 trials) for nose clip on condition and 80-100% correct (4 or 5 corrects / 5 trials) for nose clip off condition in line with previous studies (Renner et al., 2009).

### Pre-test (Smell test ortho nasal)

To check participants ortho nasal olfactory performance and to exclude anosmia we used the coffee smell test (Humphries & Singh, 2018) reported to have excellent validity with sensitivity of 93% and specificity of 96% in comparison to a 12 item Sniffin Sticks test kit (Singh et al., 2018). We prepared a 100ml cup with grounded coffee beans and one empty cup. In a trial, the participants were asked to close their eyes and sniff from a cup that was presented to them (either coffee or empty) they had to report the smell by marking on 0 – 10 scales the smell intensity. 0 indicated no smell at all with 10 indicating a very strong smell. Each participant performed 5 trials with nose clip on and 5 trials with nose clip off.

### Stimuli for the scan

For the fMRI scan the sucrose used was >99.7% pure with less than 0.04 % inverted sugar (i.e. fructose and glucose) and less than 0.06 % loss during drying, and sourced from Wiener Zucker, Feinkristallzucker, Austria and the sweet concentration of sucrose was 6% (Wee et al., 2018). Sucrose was diluted and delivered in distilled water (6g in 100ml). A tasteless control solution (containing the main ionic components of saliva, 25 mMKCl + 2.5 mMNaHCO3) was used as a rinse condition on each trial.

### Nose clips

Soft plastic foam nose clips were used to block retro nasal smell (size approx. 6.8 x 4 cm/ 2.7 x 1.6 inches in length and width) and sourced from Frienda Ltd., China). The pleasantness, pain and comfort of the nose clips was piloted before the study, on 8 subjects. All participants rated the nose clip on the nose between 4 and-4 for pleasure, pain and comfort, once at baseline and again after wearing the nose-clip for 4 minutes (the length of time they would be wearing the nose clip in each condition in the scanner).

To examine the effects of the nose clip on subjective ratings we we used a repeated measures ANOVA with ratings (3 levels, pleasantess, pain and comfort) as one within subject factor and condition (2 levels, time1 and time2) as a second within subject factor. We found no main effect of ratings (F=0.16 (2,14) p=0.85) or time (F=0.1 (1,7) p=0.75) and no ratings * time interaction (F=2 (2,14) p=0.17) (Table 1).

**Table 1:** Baseline.

| **Table 1: Baseline** |  |  | **After 4 min** |  |  |
| --- | --- | --- | --- | --- | --- |
| **Mean (SD)** |  |  | **Mean (SD)** |  |  |
| Pleas | Pain | Comfort | Pleas | Pain | Comfort |
| -0.16 | -0.76 | -0.26 | -0.21 | -0.42 | -0.78 |
| (1.09) | (1.90) | (1.51) | (1.41) | (1.81) | (1.69) |

### Study design

The fMRI scans took place at the Centre of Integrative Neuroscience and Neurodynamics (CINN) at the University of Reading. If scheduled for a morning scan participants fasted overnight, if having an afternoon scan participants fasted for 3 h (no food, only water) before the scan. 10 participants had a morning scan, and 24 participants had an afternoon scan. 60-90 minutes before scanning all the participants were given a standardized meal similar to previous studies (a banana, a cup of orange juice, 2 crackers, ∼261 total calories) with the instruction to “eat until feeling comfortably full, without overeating” similar to our previous study (Thomas et al., 2015). We asked participants to rate their hunger and mood, before the scan, on a visual analogue scale from 0 being not at all to 10 indicating the most ever felt. Subjects were screened for potential pregnancy and metal in their body before being placed in the fMRI scanner.

### Taste delivery

Tastes were delivered to the subject via separate long (∼3 m) thin Teflon tubes with a mouthpiece (∼ 1cm in diameter) at one end, that was held by the subject comfortably between the centre of the lips. At the other end of the tubes were connected to separate reservoirs via syringes and one-way Syringe Activated Dual Check Valves (Model 14044– 5, World Precision Instruments, Inc) which allowed any stimulus to be delivered manually by the researcher at exactly the right time indicated by the programme (Murray et al., 2014) thus avoiding the delays and technical issues experienced when using computerised syringe drivers.

### fMRI Task

At the beginning of a trial, a white cross at the centre of the screen appeared for 2 s indicating the start. Then, sucrose was delivered in a 0.5 mL aliquot to the subject’s mouth, the green cross was presented at the same time on the visual display for 5 s. The instruction given to the subject was to move the tongue once as soon as a stimulus was delivered in order to distribute the solution round the mouth to activate receptors, and then to keep still until a red cross was shown, when the subject could swallow. Swallowing was 2 s, then the subject was asked to rate the ‘pleasantness’ (+2 to –2) to measure hedonic value, and asked to rate the mouth fullness (richness) of the taste in their mouth (0 to +4) to measure the sensory intensity of sucrose, on a visual analogue scale by moving a bar to the appropriate point on the scale using a button box, ratings similar to those used in previous taste/fmri studies (Rolls, 2019). Each rating period was 5 s long. After the last rating on each trial 0.5mL of the tasteless control solution was administered in the same way as the sucrose stimulus at the same time as a green cross was presented on the visual display for 5 s. The control tasteless rinse was used as the comparison condition to allow somatosensory effects produced by liquid in the mouth, and the single tongue movement made to distribute the liquid throughout the mouth, to be subtracted in analysis (De Araujo et al., 2003; O’Doherty et al., 2001b). The control tasteless rinse was not subjectively rated. Then, a grey cross was presented for a duration between 0.8 s and 2 s (jittered) to indicate the end of the trial. Then the screen was black for 2 s before a new trial started. Each trial lasted ∼30 sec. Using a block design there were 7 trials of sucrose and control condition with the nose clip off. Then the scanner was stopped ∼7 to 10min, and the participant had a break before the nose clip was placed on the nose. During the break participants were told to let go of the taste tubes and just relax and they could close their eyes. Although we have shown previously no habituation effects in the subjective pleasantness and mouth fullness of sucrose after 10 presentations over the course of a 30 min task (Ko et al., 2025) we also introduced a break between the blocks in this study to avoid habituation effects and time to introduce the nose clip. After the break we ran another localiser scan followed by 7 trials of sucrose taste and control condition with nose clip on. The whole task took ∼30 minutes, including stopping and starting the scanner.

### fMRI data acquisition

Blood oxygenation level dependent (BOLD) functional MRI images were acquired using a three-Tesla Siemens scanner (Siemens AG, Erlangen, Germany) with a 32-channel head coil. During the task, around 1500 volumes were obtained for each participant, using a multiband sequence with GRAPPA and an acceleration factor of 6. Other sequence parameters included a repetition time (TR) of 700ms, an echo time (TE) of 30ms, and a flip angle (FA) of 90°. The field of view (FOV) covered the whole brain with a voxel resolution of 2.4 x 2.4 x 2.4mm^3^. Moreover, structural T1-weighted images were acquired utilizing a magnetization prepared rapid acquisition gradient echo sequence (TR = 2020ms, TE = 3.02ms, FA = 9°) with a FOV covering the whole brain and a voxel resolution of 1 x 1 x 1mm^3^.

### fMRI data analysis

The imaging data were analysed using SPM12 (Wellcome Centre for Human Neuroimaging, University College London). Pre-processing of the data used SPM12 realignment, coregister, segment, normalization to the MNI coordinate system (Montreal Neurological Institute; Collins et al., 1994), and spatial smoothing with a 6 mm full width at half maximum isotropic Gaussian kernel. The time series at each voxel was low-pass filtered with a haemodynamic response kernel. Time series non-sphericity at each voxel was estimated and corrected for, with a high-pass filter with cut-off period of 128 s.

In the single-event design, a general linear model was then applied to the time course of activation in which stimulus onsets were modelled as single impulse response functions and then convolved with the canonical hemodynamic response function. Linear contrasts were defined to test specific effects. Time derivatives were included in the basis functions set. Following smoothness estimation, linear contrasts of parameter estimates were defined to test the specific effects of each condition with each individual dataset. Voxel values for each contrast resulted in a statistical parametric map of the corresponding t statistic (transformed into the unit normal distribution (SPM z)). Movement parameters and were added as additional regressors.

At the second level, we report the main effects of sucrose with nose clip off vs the corresponding control tasteless conditions with nose clip off (supplemental data), and sucrose with nose clip on vs sucrose with nose clip off, thresholded at p<0.05 corrected (familywise-error (FWE) and p values cluster corrected at both p<0.05 False Discovery Rate (FDR) and p<0.05 FWE. We also added gender, hunger level and scan time as covariates of no interest.

We then examined regions of interest (ROI) spheres (10mm) for the anterior insula (primary taste cortex, [-32, 16, 2]) posterior insula [-38,-2,-12] and postcentral gyrus [60-16 24] using WFU pickatlas, identified in the meta-analysis on sweet tastes in humans (Roberts et al., 2020). We examined the olfactory regions; the piriform cortex, olfactory cortex and the orbitofrontal cortex using aal atlas anatomical masks in WFU pickatlas. Given our interest in retro nasal effects (Small et al., 2005) and attention to odors (Veldhuizen & Small, 2011) we also created a sphere (10mm) in the pgACC [3,42,-9] (Small et al., 2005) and examined anatomical masks of the mOFC (Small et al., 2005) and sgACC, (BA25) (Veldhuizen & Small, 2011) using aal atlas in WFU pickatlas. Finally, as we are interested in the retro nasal contribution to the rewarding effects of taste we also examined the nucleus accumbens (Berridge, 2009) and amygdala (Gottfried et al., 2003) using (IBASPM71 atlas) and aal atlas anatomical masks, respectively, in WFU pickatlas. Data were extracted using the SPM ROI analysis Matlab code and MarsBar and analysed with paired-sample t tests in excel. We also examined correlations between the extracted ROI data and the subjective ratings of pleasantness and mouth fullness. As we are also interested in the effects of retro nasal processing on the timing of activity, we also examined the time to peak activity within 10 s, in the nose clip on vs off conditions. This was examined by calculating the time to maximum peak activity within the first 10 s after the onset of sucrose delivery from the time course data, using the max function in Matlab.

## Results

### Demographic data for fMRI study

34 participants took part with a mean age of 25 yrs. See Table 2 for demographics.

**Table 2:**
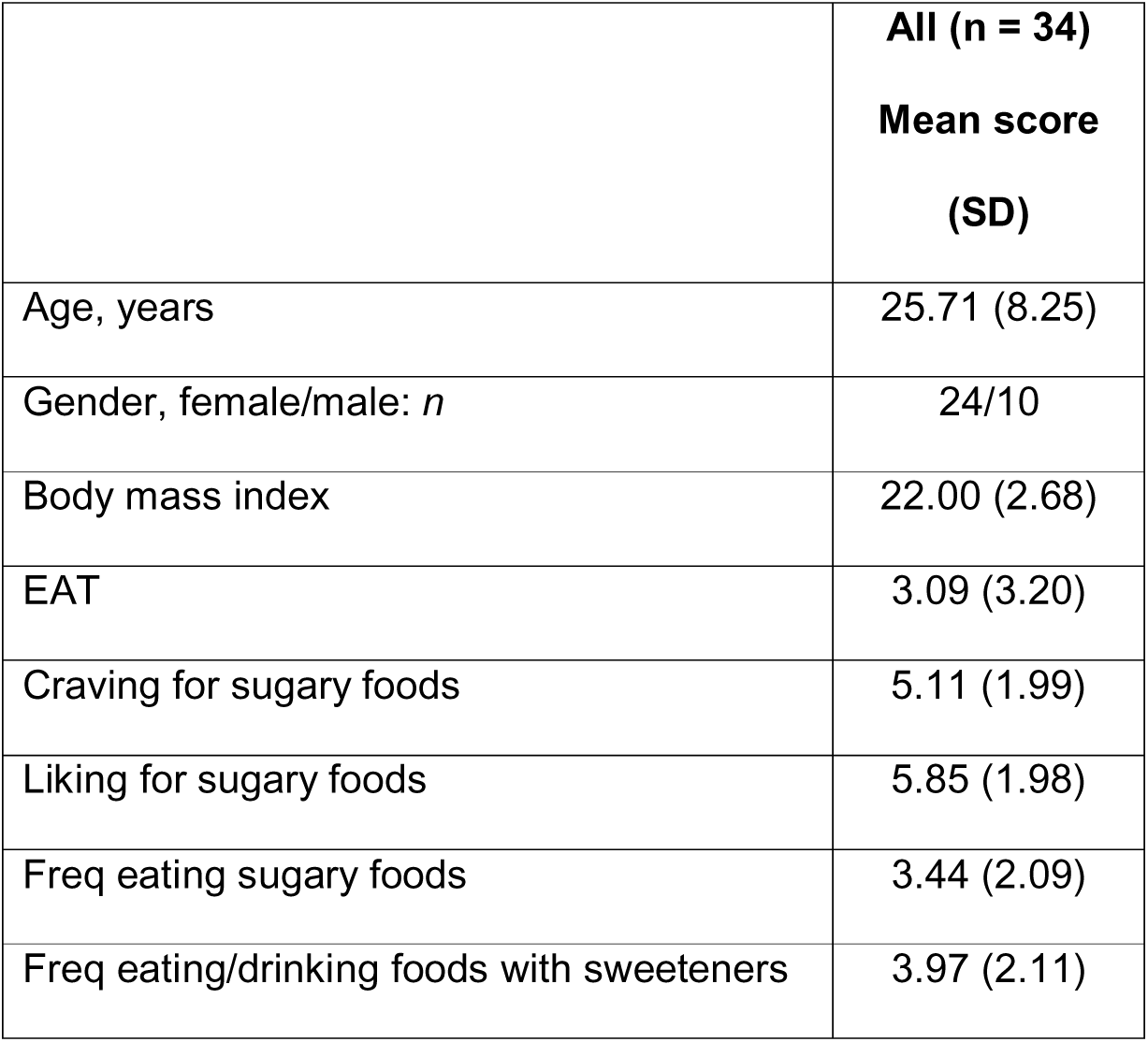
Demographics.

| **Table 2: Demographics** | **All (n = 34)**<br>**Mean score**<br>**(SD)** |
| --- | --- |
| Age, years | 25.71 (8.25) |
| Gender, female/male: *n* | 24/10 |
| Body mass index | 22.00 (2.68) |
| EAT | 3.09 (3.20) |
| Craving for sugary foods | 5.11 (1.99) |
| Liking for sugary foods | 5.85 (1.98) |
| Freq eating sugary foods | 3.44 (2.09) |
| Freq eating/drinking foods with sweeteners | 3.97 (2.11) |

### Pre-test results of sensitivity to 2% sucrose

Twenty-one participants passed the pre-test with 6 out of 6 trials correct the first time. Ten participants passed the pre-test with 5 out of 6 trials correct the first time and three participants got 6 of the 6 trials correct on their second attempt, so were also included in the study.

### Pre-test Candy Smell Test

As expected we found that with the nose clip off participants could identify the flavours in the candy smell test with average accuracy of 84% (± 14) and when the nose clip was added this accuracy dropped to 31% (± 20) in line with previous studies (Renner et al., 2009).

### Pre-test (Smell test ortho nasal)

As expected, all participants could identify the cup that had coffee in compared to no coffee and rated the coffee smell as above average intensity and rated the coffee smell (6.70 ±1.78) higher than the intensity of the empty cup smell (1.17 ±1.90), (t(22) = 12.04, p <0.001) and rated the intensity of the coffee smell (6.70 ±1.78) higher with the nose clip off than with the nose clip on (0.26 ±0.59), (t(22) = 16.7, p <0.001).

### fMRI scan day

#### Subjective hunger and Mood

Participants had relatively high mood and low hunger levels before the scan (Table 3).

**Table 3:**
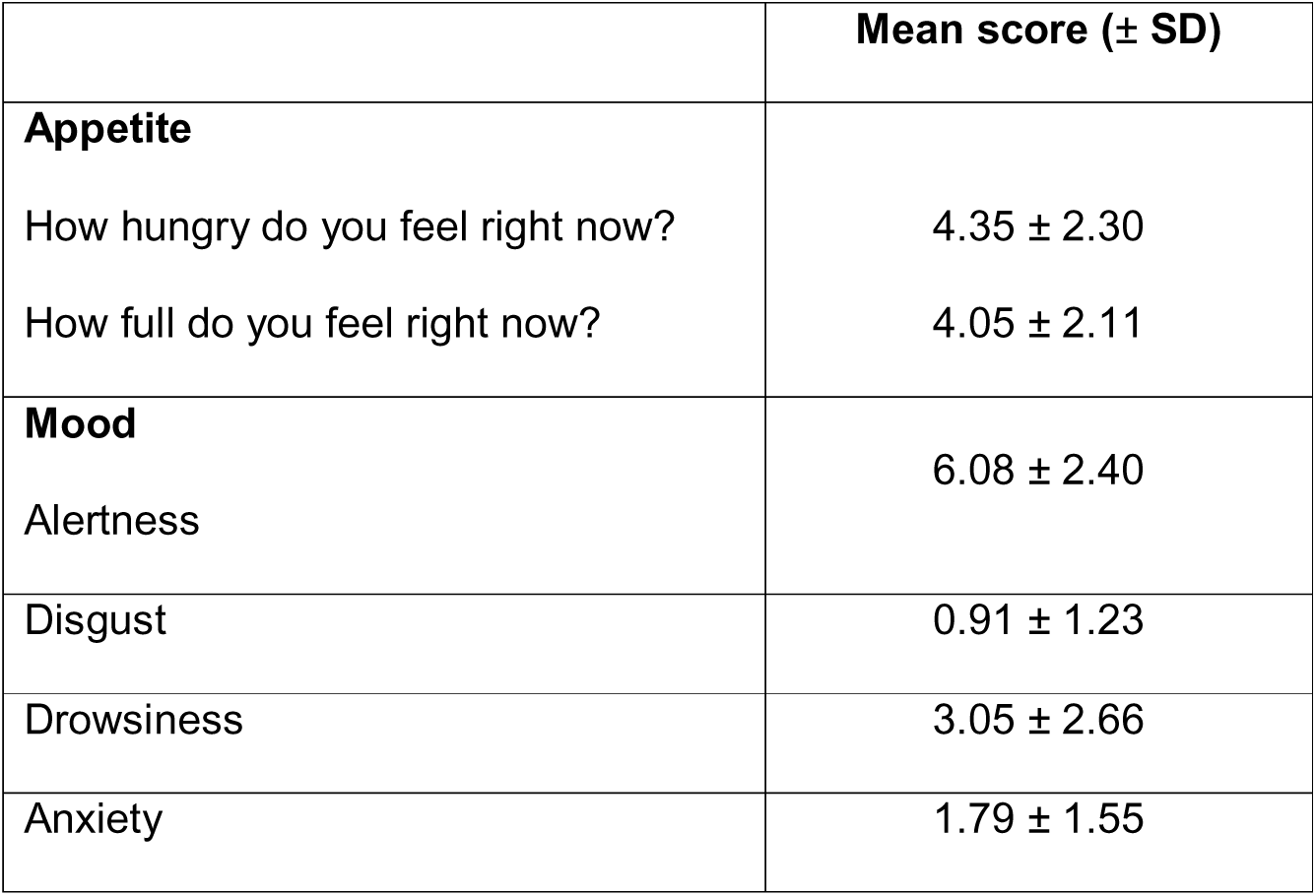
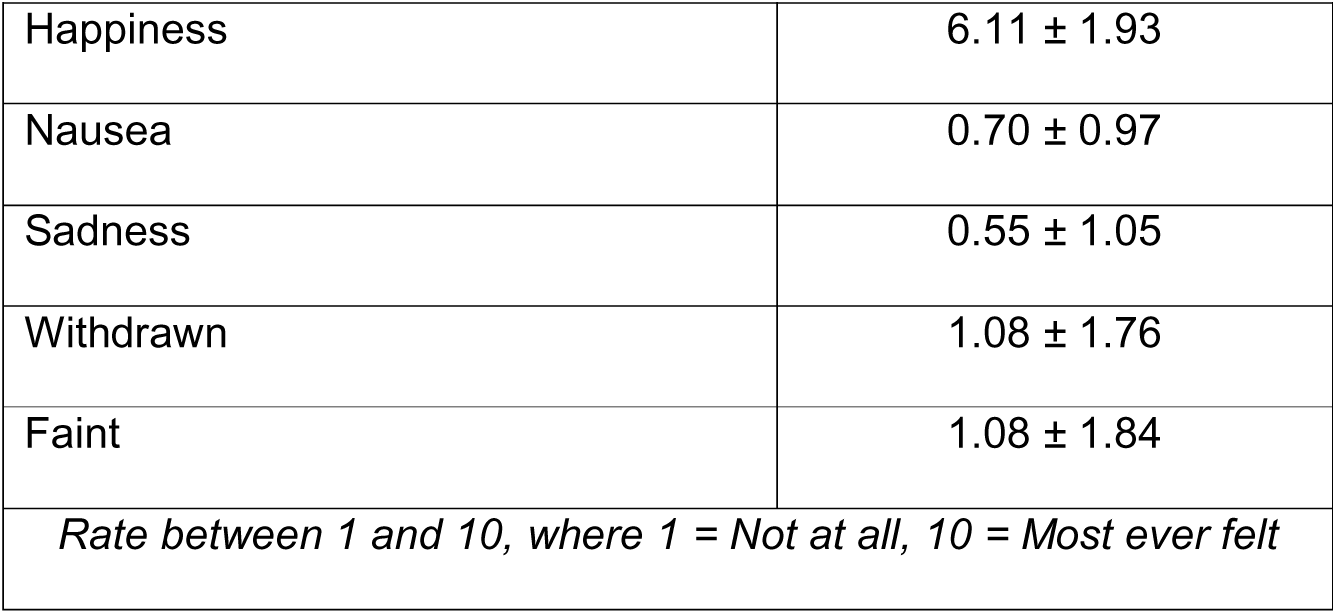
Visual Analogue Scale.

| **Table 3: Visual Analogue Scale** | **Mean score (± SD)** |
| --- | --- |
| **Appetite** |  |
| How hungry do you feel right now? | $4.35 \pm 2.30$ |
| How full do you feel right now? | $4.05 \pm 2.11$ |
| **Mood** |  |
| Alertness | $6.08 \pm 2.40$ |
| Disgust | $0.91 \pm 1.23$ |
| Drowsiness | $3.05 \pm 2.66$ |
| Anxiety | $1.79 \pm 1.55$ |
| Happiness | 6.11 ± 1.93 |
| Nausea | 0.70 ± 0.97 |
| Sadness | 0.55 ± 1.05 |
| Withdrawn | 1.08 ± 1.76 |
| Faint | 1.08 ± 1.84 |
| *Rate between 1 and 10, where 1 = Not at all, 10 = Most ever felt* |  |

#### Subjective ratings of stimuli: Pleasantness and Fullness ratings during the scan with nose clip on and off

To check for any habituation effects we examined the pleasantness and fullness ratings at the beginning and the end of block 1, trial 1 vs trial 7, the nose clip off condition, using paired samples t-test. We found no significant differences in pleasantness t(32) = 0.65, p = 0.51, or mouth fullness t(31) = 1.88, p =0.07 from the beginning to the end of the block, as expected, suggesting no habituation to sucrose taste over the trials.

To examine the effects of the nose clip on subjective ratings we we used a repeated measures ANOVA with ratings (2 levels, pleasantness, mouth fullness) as one within subject factor and condition (2 levels, nose clip on, nose clip off) as a second within subject factor. We found a main effect of ratings (F=27.5 (1,33) p<0.001) and a main effect of condition (F=6.6 (1,33) p=0.015) but no ratings * condition interaction (F=0.39 (1,33) p=0.54) (Figure 1). Follow up paired sample t-tests showed that mouth fullness was rated higher for nose clip off than nose clip on t(33) = 2.5, p =0.017.

**Figure 1.**
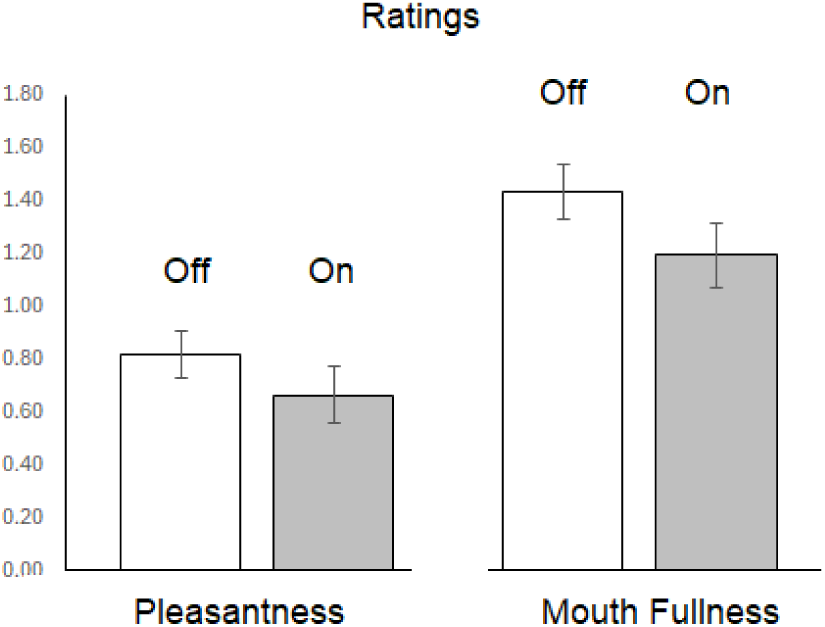
Pleasantness and Mouth Fullness rating for sucrose nose clip on and nose clip off conditions in the scanner.

**Figure 2.**
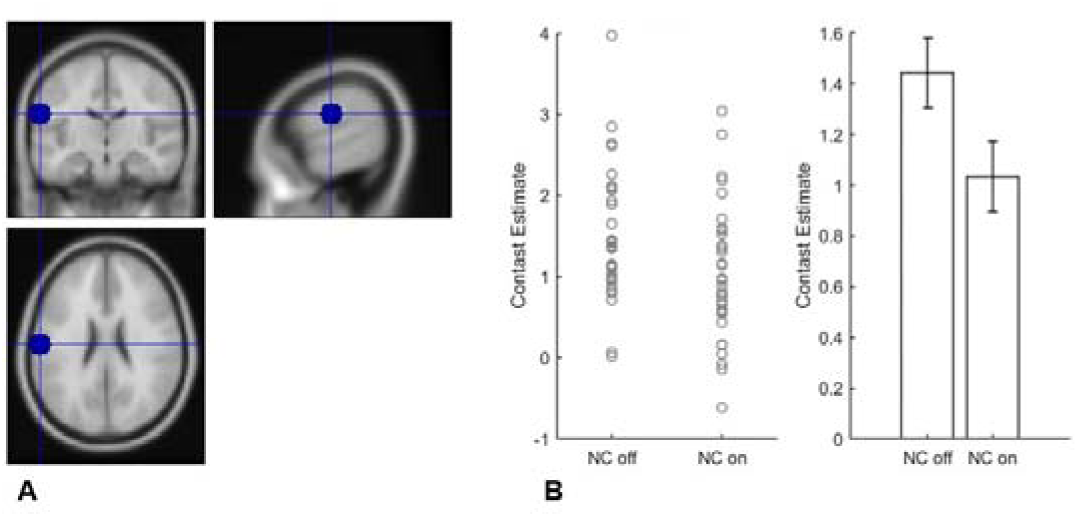
A. Left postcentral gyrus ROI. B. Contrast estimates extracted from ROI using marsbar for sucrose nose clip off and nose clip on conditions, error bars SEM.

**Figure 3.**
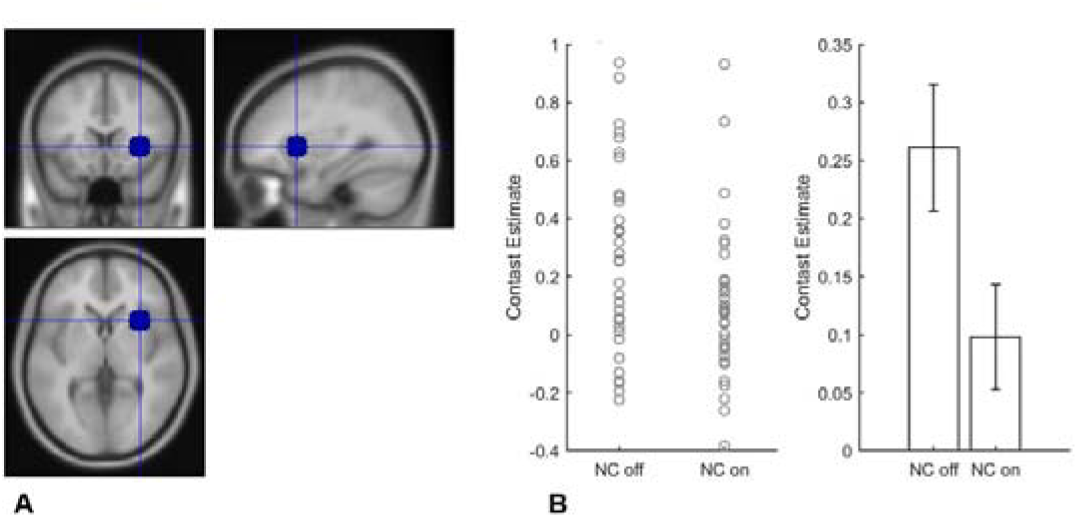
A. Right Anterior Insula ROI. B. Contrast estimates extracted from ROI using marsbar for sucrose nose clip off and nose clip on conditions, error bars SEM.

**Figure 4.**
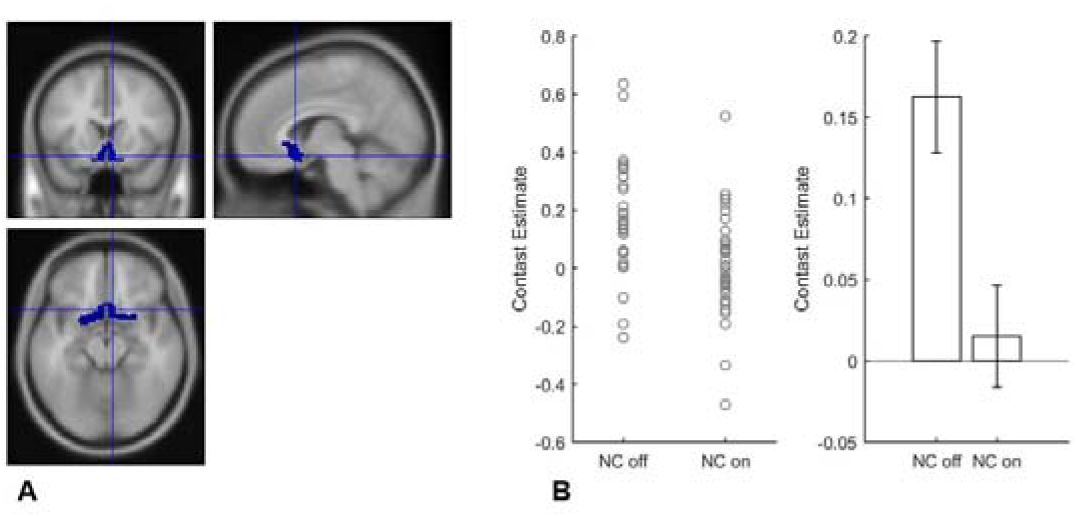
A. Olfactory Cortex ROI. B. Contrast estimates extracted from ROI using marsbar for sucrose nose clip off and nose clip on conditions, error bars SEM.

**Figure 5.**
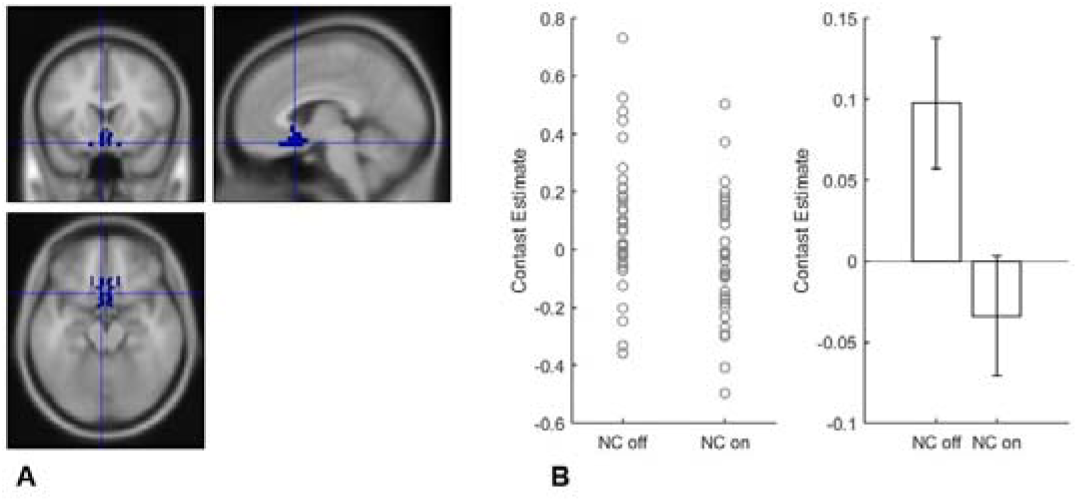
A. sgACC (BA25) ROI. B. Contrast estimates extracted from ROI using marsbar for sucrose nose clip off and nose clip on conditions, error bars SEM.

**Figure 6.**
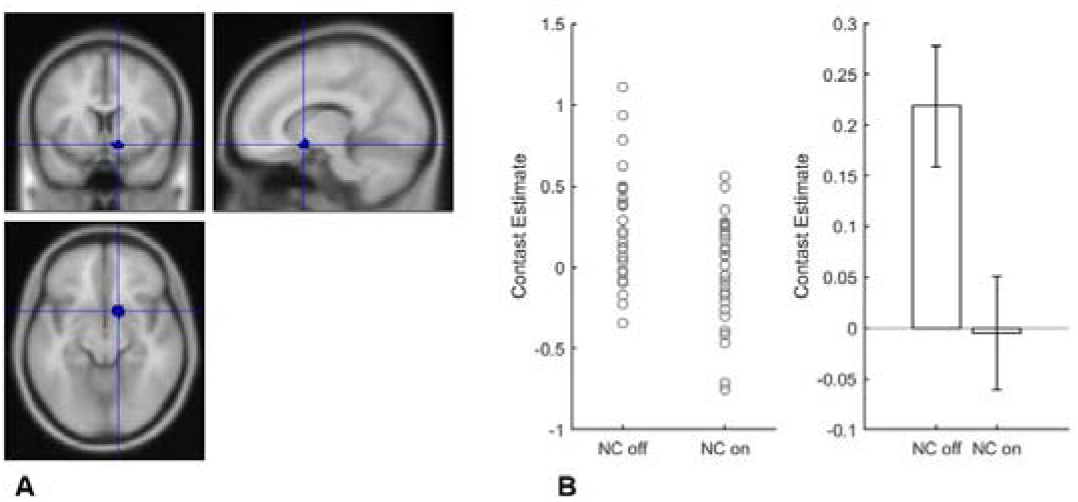
A. Right NAcc ROI. B. Contrast estimates extracted from ROI using marsbar for sucrose nose clip off and nose clip on conditions, error bars SEM.

### ROI analysis

#### Nose clip off vs nose clip on

We found greater neural activity in the sucrose nose clip off condition vs the sucrose nose clip on condition (Table 4). The right postcentral gyrus, right anterior and right posterior insula, the olfactory cortex, piriform cortex, sgACC and right NAcc activity survived when controlling for multiple comparisons (Table 4).

**Table 4.**
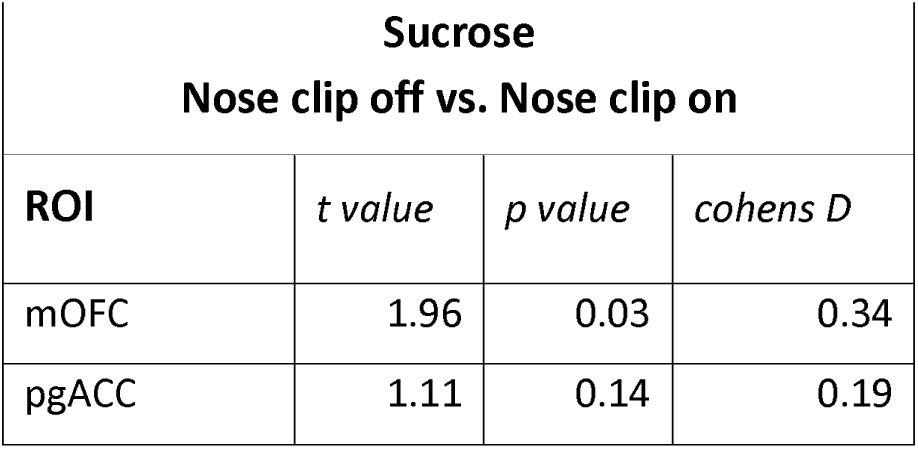
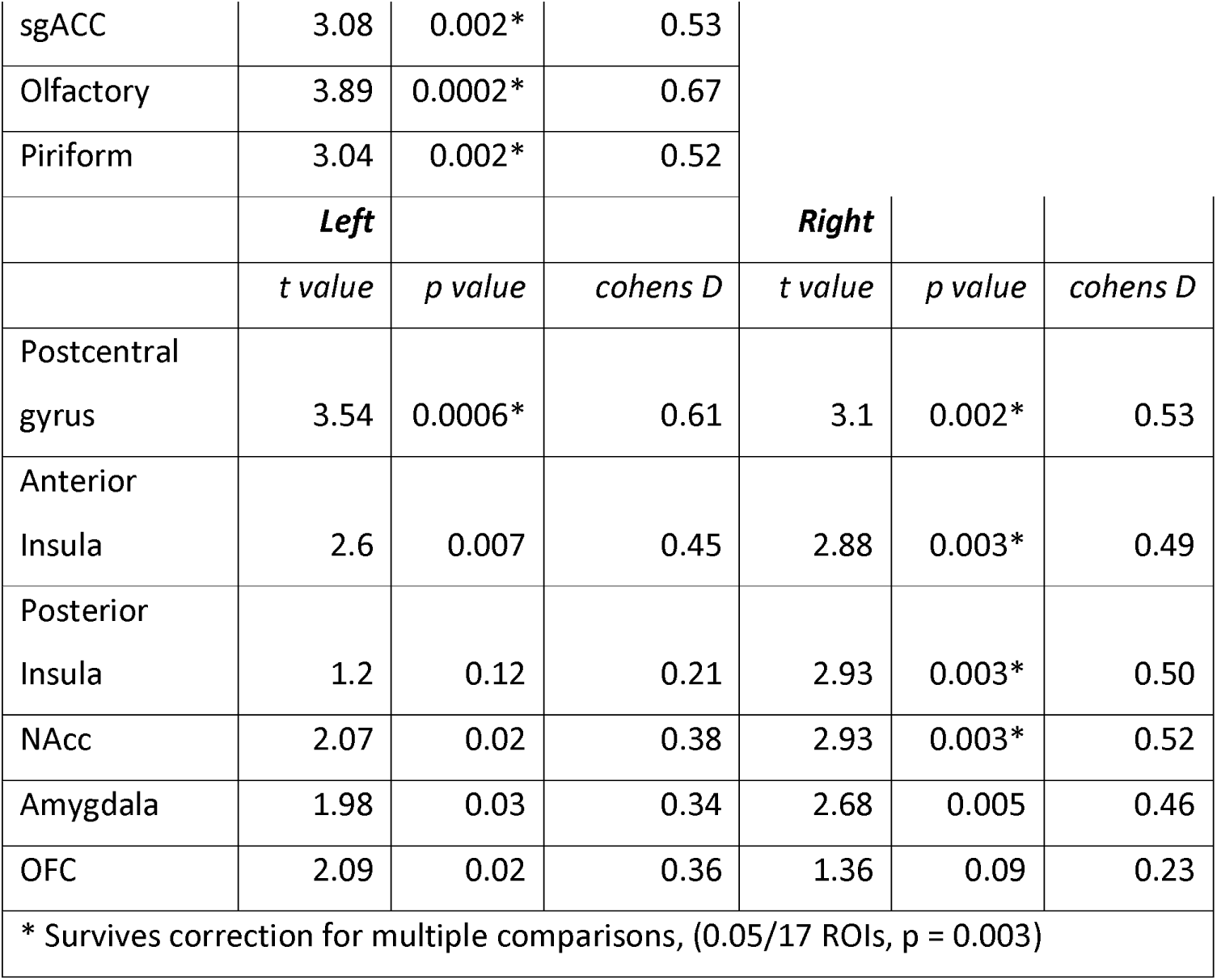
ROI Results.

### Temporal effects

We also examined the time to peak activity after the taste of sucrose in the nose clip on vs off conditions. Using repeated measures ANOVA with ROIs as one within subject factor and condition (nose clip on, nose clip off) as a second within subject factor. We found a main effect of ROI (F=12.6 (1,8) p<0.001) but no main effect of condition and no ROI * condition interaction (Figure 7).

**Figure 7.**
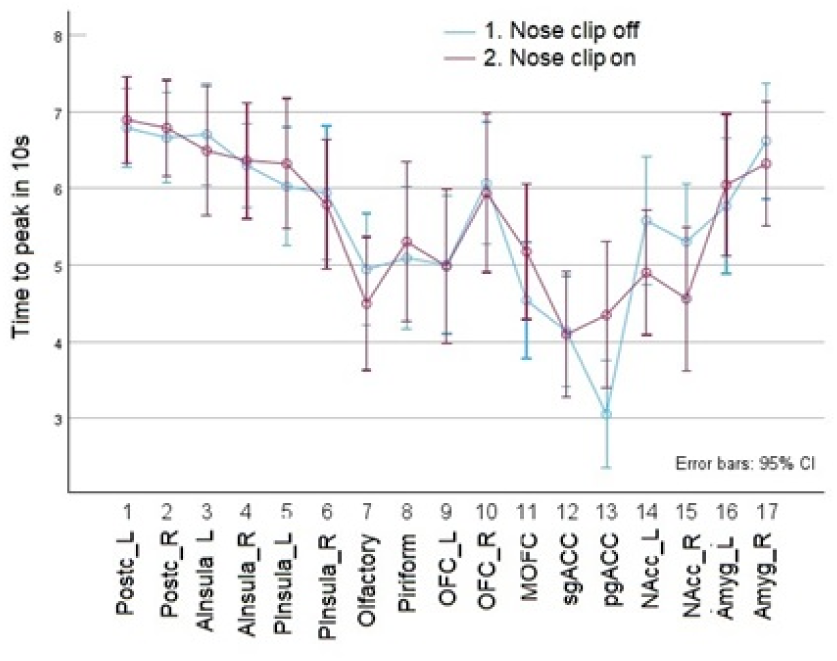
Time to first peak (10 s time bin) for each ROI and for each condition: nose clip on and nose clip off.

**Figure 8.**
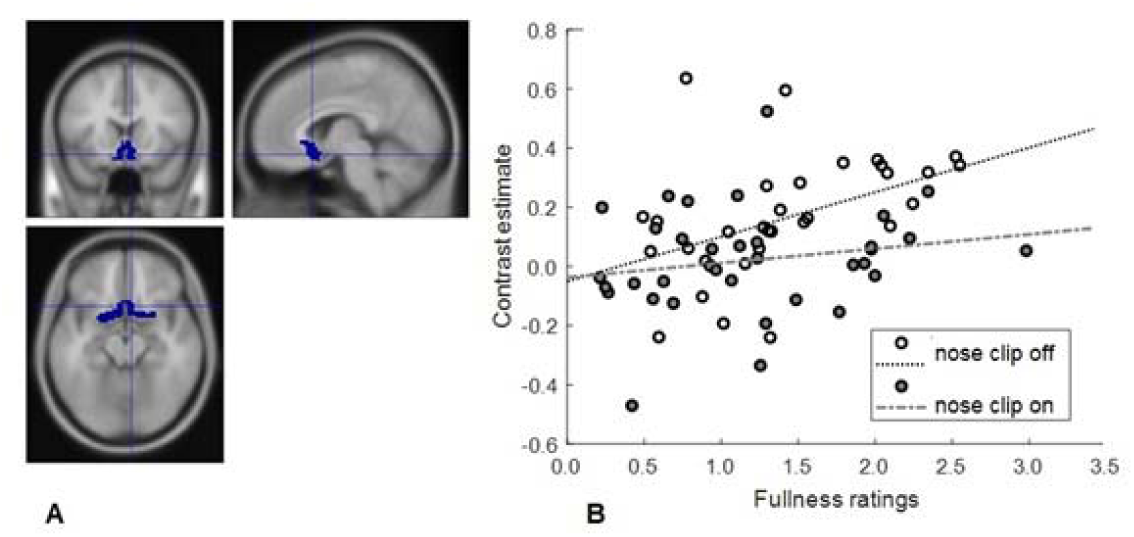
A. Olfactory cortex ROI. B. Correlations between mouth fullness ratings and brain activity to sucrose with the nose clip on and nose clip off conditions.

### Parametric Modulation

We also found positive correlations between mouth fullness ratings and activity in ROIs olfactory cortex (rho1 = 0.44, p = 0.01), the sgACC (rho1 = 0.34, p = 0.046), the pgACC (rho1 = 0.37, p =0.03) and the mOFC (rho1 = 0.36, p = 0.036) for nose clip off, but these did not survive correction for multiple comparisons. No correlations between ratings and ROI data were found for the nose clip on condition.

### Exploratory Whole brain analyses

#### Main effects of taste stimuli

The sucrose taste vs the control taste activated regions such as the primary taste cortex (insula), primary somatosensory cortex (postcentral gyrus), and the precentral gyrus and caudate, similar to previous studies on sweet tastes (Yeung & Wong, 2020) (Table S1). There were no significant activations for the opposite contrast, control taste vs sucrose taste.

#### Nose clip off vs on

When examining the whole brain results (Table 5) we found reduced activity for the contrast sucrose nose clip off vs sucrose nose clip on in regions such as the rolandic operculum, precuneus and post central gyrus, these results were apparent only when using a p=0.001 uncorrected threshold. There were no regions activated under the contrast sucrose nose clip on vs sucrose nose clip off, at any threshold. There was reduced precuneus activity for the contrast tasteless control nose clip off vs tasteless control nose clip on at p=0.001 uncorrected threshold. There were no regions activated under the contrast tasteless control nose clip on vs tasteless control nose clip off, at any threshold.

**Table 5.**

| **Table 5:**<br>**Sucrose**<br>**(Nose Clip Off – Nose Clip On)**<br>*Threshold: $p=0.001$ uncorrected* |  |  |  |  |  |  |  |
| --- | --- | --- | --- | --- | --- | --- | --- |
| Region | x(mm) | y(mm) | z(mm) | Z-score | Num of voxel | p(FWE-corr) | p(FDR-corr) |
| Rolandic operculum | 40 | 1 | 16 | 5.07 | 71 | =0.036 | =0.060 |
| Precuneus | -1 | -83 | 43 | 4.82 | 143 | =0.001 | =0.003 |
| Lingual gyrus | -16 | -64 | 0 | 4.18 | 227 | < 0.0001 | <0.001 |
| Postcentral gyrus | -61 | -26 | 24 | 4.08 | 146 | =0.001 | =0.003 |
| Rolandic operculum | 56 | 3 | 7 | 4.06 | 91 | =0.013 | =0.026 |

## Discussion

This is the first study to examine both the subjective experience and the objective brain response to sucrose taste in humans when olfactory processing is occluded with a nose clip. In support of our hypothesis, we provide novel evidence that a nose clip reduces the subjective pleasantness of sucrose taste. These results are consistent with previous studies finding reduced sucrose sweetness with a nose clip on (Mu et al., 2024; Yang et al., 2024). We also extend the previous literature by showing that a nose clip reduces the subjective mouth fullness of the sucrose taste. Together this work supports the view that retro nasal pathways may contribute to the perceptual processing of “non-volatile” substances such as sucrose (He et al., 2024; He et al., 2023).

We know of no studies examining the effects on the human brain of occluding retro nasal olfactory processing when tasting sucrose. We provide novel findings that tasting sucrose with a nose clip results in reduced brain activity in the primary taste and olfactory regions. Specifically, we found that the postcentral gyrus, part of the somatosensory cortex modulated by sweet taste intensity (van Meer et al., 2023) the anterior and posterior insula (Dalenberg et al., 2015; Roberts et al., 2020) and the olfactory cortex and piriform cortex, had reduced activity when tasting sucrose with a nose clip on. We also found that brain activity tracked subjective mouth fullness with the nose clip off but not with nose clip on. Together these data further support the view that retro nasal pathways could contribute to sucrose taste processing (He et al., 2024; He et al., 2023).

Interestingly we found the secondary olfactory areas (OFC) less impacted by occlusion with the nose clip. This could be because the OFC as a multimodal region is much less dependent on signals coming purely from one modality (Rolls, 2019) and therefore is still activated by the taste in the mouth with the nose clip on. This also fits with the study by Yang et al where the nose clip affected the processing of tastes and aromas but not other auditory or visual senses (Yang et al., 2024).

We also found that the nose clip reduced neural activity in the sgACC, a region that plays a role in attention to odours (Sabri et al., 2005; Veldhuizen & Small, 2011). Although in this study participants were not asked explicitly to attend or ignore the stimulus, they were likely involuntarily attending as they were required to make subjective ratings of pleasantness and mouth fullness, on each trial. We also found that the sgACC tracked the subjective mouth fullness of the sucrose with the nose clip off but not with nose clip on, similar to that seen in the olfactory cortex mentioned above and the neighbouring pgACC and mOFC ROIs. These prefrontal cortex multi-modal regions have also been shown to have greater activation to taste when combined with a savory odour than to the sum of the activations by the taste and olfactory components presented separately (McCabe & Rolls, 2007; Rolls, 2019). Together these findings suggest that the nose clip reduces the integration of taste and olfactory components of the sucrose taste making it more difficult to perceive.

When examining the effects of the nose clip on vs off we also found reduced activity in the NAcc. The NAcc is a crucial hub in the brain related to feeding behaviour, integrating signals from both homeostatic and hedonic circuits, to facilitate behavioural output via its downstream projections (Marinescu & Labouesse, 2024). Previous studies show that the ventral striatum, of which the NAcc is part, is at the crossroads of olfactory and reward pathways and receives direct projections from the primary olfactory cortex (Ubeda-Bañon et al., 2007) and the dopaminergic midbrain (Ikemoto, 2007) and is greatly involved in odour-guided eating behaviour (Murata, 2020). Hence reduced activity in this region with the nose clip on supports the idea that potential retro nasal olfactory signals from the sucrose taste could be being occluded.

When examining the exploratory whole brain data, we found that the nose clip occlusion reduced neural activity to sucrose taste in the rolandic operculum and precuneus. The rolandic operculum plays a central role in flavour percept formation (Small et al., 2004) with recent studies showing that neural taste and smell signals are integrated in the rolandic operculum (Suen et al., 2021).The operculum, is a large structure with three lobes and a complex array of functions including sensory, motor, autonomic and cognitive processing. In humans, these are extended with the addition of language (Mălîia et al., 2018). Studies mapping the function of the rolandic operculum, using direct electrical stimulation, find it involved in oropharyngeal responses with the most widespread and common mapping it to the pharynx–larynx or the tongue (Mălîia et al., 2018). Further when stimulated participants report experiencing taste, making the rolandic operculum a likely candidate for the primary gustatory cortex (Mălîia et al., 2018). Connections between the rolandic operculum and the anterior and posterior insula support its role in feeding behaviour while connections with the frontal operculum, premotor area, fusiform gyrus and post central gyrus support its role in speech production (Mălîia et al., 2018). Due to such connections, some suggest a link between flavour perception and language development, citing gustation-language connectivity and chimpanzees’ vocal food communications (Kalan et al., 2015; Schel et al., 2013).

Finally, we also found reduced precuneus activity to sucrose taste with the nose clip on. The precuneus is primarily involved in complex cognitive functions like episodic memory retrieval, self-processing, visuo-spatial imagery, and imagining future events, essentially acting as a key hub for integrating personal experiences and constructing mental scenarios; it is considered a core part of the brain’s “default mode network” which is active during resting states and internal thought processes (Cavanna & Trimble, 2006). Therefore, reduced activity in this region to sucrose taste (and also to the tasteless control) with the occlusion of the retro nasal pathways could reflect a difficulty in determining the percept of the stimuli and a need to recruit memories of previous taste experiences.

Taken together, we provide neuroscientific evidence of how retro nasal olfaction could contribute to taste perception by influencing brain activity in taste, smell, attention, and reward regions. These findings suggest that the view of “non-volatile” sucrose taste alone in oral cavity does not fully explain the reduced subjective and neural effects of sucrose with the nose clip on. We suggest that perhaps retro nasal sensations are playing a role in sucrose perception.

In conclusion, knowing how the retro nasal pathway blockade may impact basic taste processing could help explain how individuals with olfactory impairments (e.g., due to aging, illness, or COVID-19) experience reduced appetite and altered eating behaviours. Further studies are required to test if modifying sensory perception of sweet tastes with aromas could help the design of more satisfying low-sugar foods. In sum, this study contributes to a broader understanding of how sensory processing shapes eating experiences, with applications in neuroscience, health, and food innovation.

## Acknowledgements

We would like to thank Dr Shan Shen and the staff at the Centre for Neuroscience and Neurodynamics (CINN) at the University of Reading for their help with the scanning. We would also like to thank Dr. Jing-guo Chen, The Second Affiliated Hospital of Xi’an Jiaotong University, Xi’an, China for discussion on Brain Regions Involved in Taste and the Physiology of the Retronasal Cavity and Dr. Bao-qing Zhu, Beijing Forestry Unversity, Beijing, China for discussion on sensory evaluation.

## Author Contributions

CMcC, TE, JS, WS conceived and designed research. JS, WS and TE supplied the sucrose. HK and TE conducted research and analysed the data supported by CMcC. CMcC and HK drafted the article. All authors actively participated in editing and reviewing the manuscript.

## Data Accessibility Statement

The data that support the findings of this study are available from the corresponding author, upon reasonable request.

