## Supplement for "Retro Nasal blockade reduces the Neural Processing of Sucrose in the Human Brain"

*Main effects of taste stimuli*

**Table S1**

| **Sucrose - control (Nose Clip Off)**  Threshold: p=0.05 FWE corrected | | | | | | | |
| --- | --- | --- | --- | --- | --- | --- | --- |
| **Region** | **x(mm)** | **y(mm)** | **z(mm)** | **Z-score** | **voxels** | **p(FWE-corr)** | **p(FDR-corr)** |
| Postcentral | -42 | -18 | 52 | 6.314 | 152 | < 0.0001 | < 0.0001 |
| Precentral | -35 | -23 | 55 | 6.022 |  |  |  |
| Supp Motor Area | 4 | 10 | 55 | 6.059 | 252 | 0 | < 0.0001 |
| Supp Motor Area | -6 | 15 | 48 | 5.877 |  |  |  |
| Insula | 37 | 20 | 7 | 5.704 | 106 | < 0.0001 | < 0.0001 |
| Mid Cingulate gyrus | -8 | 25 | 31 | 5.466 | 10 | =0.000863 | =0.050467 |
| Hippocampus | 42 | -30 | -8 | 5.400 | 22 | < 0.0001 | =0.00449 |
| Insula | -35 | 15 | 7 | 5.260 | 29 | < 0.0001 | =0.001665 |
| Caudate | 18 | 15 | 7 | 5.131 | 23 | < 0.0001 | =0.004377 |
| Insula | -42 | 18 | 0 | 5.131 | 10 | 0.000863 | 0.050467 |
| Threshold: 0.0001 uncorrected | | | | | | | |
| **Region** | x(mm) | y(mm) | z(mm) | Z-score | voxels | p(FWE-corr) | p(FDR-corr) |
| Superior frontal gyrus | -23 | -4 | 55 | 5.082 | 518 | < 0.0001 | < 0.0001 |
| Inferior parietal | -54 | -23 | 48 | 3.736 |  |  |  |
| Insula | 37 | 20 | 7 | 5.704 | 1185 | 0 | < 0.0001 |
| Putamen | 23 | 15 | -8 | 5.244 |  |  |  |
| Frontal operculum | 47 | 3 | 24 | 4.822 |  |  |  |
| Hippocampus | 42 | -30 | -8 | 5.400 | 160 | < 0.0001 | < 0.0001 |
| Superior temporal gyrus | 37 | -33 | 7 | 4.938 |  |  |  |
| Cerebellum | 20 | -59 | -22 | 4.943 | 28 | =0.02442 | =0.049727 |
| Insula/Rolandic operculum | 37 | -4 | 14 | 4.911 | 31 | =0.018074 | =0.039507 |
| Superior parietal | -28 | -57 | 57 | 4.894 | 74 | =0.000476 | =0.001917 |
| Precuneus | -13 | -59 | 64 | 3.787 |  |  |  |
| Mid frontal cortex | 40 | 34 | 19 | 4.561 | 39 | =0.008436 | =0.01988 |

**Table S2**

| **Sucrose - control (Nose Clip On)**  Threshold: p=0.05 FWE corrected | | | | | | | |
| --- | --- | --- | --- | --- | --- | --- | --- |
| **Region** | **x(mm)** | **y(mm)** | **z(mm)** | **Z-score** | **voxels** | **p(FWE-corr)** | **p(FDR-corr)** |
| Supp Motor Area | 4 | 20 | 48 | 6.565 | 277 | 0 | < 0.0001 |
| Supp Motor Area | -6 | 13 | 50 | 6.010 |  |  |  |
| Caudate | -6 | 10 | 4 | 6.237 | 199 | < 0.0001 | < 0.0001 |
| Caudate | 11 | 15 | 7 | 6.202 |  |  |  |
| Postcentral gyrus | -37 | -21 | 52 | 6.390 |  |  |  |
| Precentral gyrus | -37 | -26 | 60 | 6.296 | 304 | 0 | < 0.0001 |
| Precentral gyrus | 52 | 6 | 33 | 5.708 | 40 | < 0.0001 | =0.000138 |
| Insula | 32 | 18 | 7 | 5.044 | 32 | < 0.0001 | =0.000486 |
| Inferior frontal gyrus | -40 | 25 | 26 | 5.229 | 17 | =0.000142 | =0.007097 |
| Inferior frontal operculum | -47 | 22 | 31 | 5.202 |  |  |  |
| Insula | -30 | 25 | 9 | 5.225 | 22 | < 0.0001 | =0.00264 |
| Threshold: 0.0001 uncorrected | | | | | | | |
| Region | x(mm) | y(mm) | z(mm) | Z-score | Num of voxel | p(FWE-corr) | p(FDR-corr) |
| Mid cingulate gyrus | 11 | 30 | 31 | 4.590 | 1022 | 0 | < 0.0001 |
| Superior anterior cingulate | -8 | 25 | 28 | 4.335 |  |  |  |
| Medial superior frontal gyrus | 11 | 32 | 43 | 4.048 |  |  |  |
| Superior parietal | 30 | -59 | 57 | 4.012 | 282 | < 0.0001 | < 0.0001 |
| Inferior parietal | 40 | -54 | 50 | 4.722 |  |  |  |
| Precuneus | 6 | -74 | 43 | 4.367 | 72 | =0.000518 | =0.002019 |
| Mid Occipital | -28 | -90 | 14 | 4.246 | 22 | =0.04496 | =0.09369 |
